# Unique role for caspase-8 in the release of IL-1β and active caspase-1 from viable human monocytes during *Toxoplasma gondii* infection

**DOI:** 10.1101/2022.12.18.520939

**Authors:** William J. Pandori, Stephanie Y. Matsuno, Tiffany H. Kao, Sharmila Mallya, Sarah N. Batarseh, Melissa B. Lodoen

**Affiliations:** Department of Molecular Biology & Biochemistry and the Institute for Immunology, University of California, Irvine, California, 92617, USA

**Author notes:** To whom correspondence should be addressed: Dr. Melissa B. Lodoen, University of California, Irvine, 3238 McGaugh Hall, Irvine, CA 92617.

## Abstract

Monocytes are among the first cells recruited to sites of infection and major producers of the potent proinflammatory cytokine IL-1β. We previously showed that IL-1β release during *Toxoplasma gondii* infection of primary human monocytes requires the NLRP3 inflammasome and caspase-1 activity but is independent of gasdermin D and pyroptosis. To investigate potential mechanisms of pyroptosis-independent release of IL-1β during *T. gondii* infection, we constructed caspases-1, -4, -5, or -8 knockout THP-1 monocytic cells. Genetic ablation of caspase-1 or -8, but not caspase-4 or -5, decreased IL-1β release during *T. gondii* infection without affecting cell death. In contrast, TNF-α and IL-6 secretion were unperturbed in caspase-8 knockout cells during *T. gondii* infection. Dual pharmacological inhibition of caspase-8 and RIPK1 in primary monocytes also decreased IL-1β release without affecting cell viability or parasite infection efficiency. In addition, caspase-8 was required for the release of active caspase-1 from *T. gondii*-infected cells and for IL-1β release during infection with the related apicomplexan parasite *Neospora caninum*. Surprisingly, caspase-8 was dispensable for the synthesis and cleavage of IL-1β, but caspase-8 deficiency resulted in the *retention* of mature IL-1β within cells. Our data indicate that during *T. gondii* infection of human monocytes, caspase-8 functions in a novel gasdermin D-independent mechanism controlling IL-1β release from viable cells. This study expands on the known molecular pathways that promote IL-1β in human immune cells and provides the first evidence of a role for caspase-8 in the mechanism of IL-1β release during host defense against infection.

## Introduction

Caspases are a set of cysteine-aspartic acid proteases that control inflammation and cell death. In humans, caspases-1, -4 and -5 are considered inflammatory caspases, as their activities lead to the cleavage and release of the proinflammatory cytokine IL-1β and other IL-1 family cytokines (1–4). Cell death-related caspases include the initiator caspases-2, −8, −9, and −10 and the executioner caspases-3, −6 and −7. Although caspase-9 is traditionally considered an initiator of intrinsic apoptosis and caspase-8 an initiator of extrinsic apoptosis, they can also be coordinately activated (5). Both caspases serve to activate caspases-3 and −7, which execute the final stages of apoptosis (6, 7). Caspase-8 also negatively regulates an inflammatory form of cell death termed necroptosis through its inhibition of RIPK1 and downstream RIPK3 and MLKL (8). Caspase-8 can also function as an inflammatory caspase by influencing IL-1β release through priming and activation of the NLRP3 inflammasome, as well as cleaving pro-IL-1β and gasdermin D, in addition to its previously described role as an initiator caspase in apoptosis (7–13).

The proinflammatory cytokine and pyrogen IL-1β is critical for protection against many bacterial, viral, fungal, and parasitic pathogens (14, 15). However, given its potency, IL-1β has also been implicated in the development of autoimmune diseases, such as rheumatoid arthritis, atherosclerosis, type II diabetes, CAPS, Alzheimer’s disease, and gout (16–19). Modified IL-1 receptor antagonist, anakinra, is approved for the treatment of rheumatoid arthritis; however, there are no approved treatments that specifically inhibit the aberrant production or release of IL-1β, reinforcing the importance of understanding the pathways that lead to the production and release of IL-1β.

Pro-IL-1β is translated as an inactive protein, and must be proteolytically processed, most commonly by caspase-1, to exert its biological effects (3, 20, 21). Caspase-1 is activated by multiprotein complexes called inflammasomes, which consist of NOD-like receptor proteins (NLRPs) that sense intracellular stimuli and bind the adaptor protein ASC, which contains a caspase activation and recruitment domain (CARD) (22, 23). Once nucleated in these large complexes, ASC binds pro-caspase-1, bringing the proteases into close physical contact and allowing for auto-proteolysis and activation of caspase-1 (24, 25). Once cleaved, IL-1β does not exit the cell through the conventional secretory pathway via the Golgi (26). Instead, caspase-1 also cleaves gasdermin D (GSDMD), which forms pores in the cell membrane and allows IL-1β to pass through these pores from still viable cells (27, 28). Eventually these pores allow for osmosis-driven cell swelling and lysis, during which intracellular contents, including additional IL-1β, are released from the dying cells in a form of inflammatory cell death called pyroptosis (1, 29, 30).

The NLRP3 inflammasome is the best studied inflammasome and responds to a wide variety of stimuli. Canonical NLRP3 inflammasome activity in most cells requires two signals. First, a priming signal, such as LPS binding to TLR4, is required to induce NF-κB activation and nuclear translocation, leading to the transcriptional activation of pro-IL-1β and NLRP3 (25). Next, an activating signal, such as exposure to extracellular ATP or nigericin, is required to induce conformational changes in the NLR, which facilitates its oligomerization and formation of the inflammasome, activating caspase-1 and cleaving IL-1β (25, 31).

Whereas murine and human macrophages predominantly regulate IL-1β via this model, human monocytes and neutrophils activate the NLRP3 inflammasome and release IL-1β in response to a single priming stimulus and do not require a secondary activating signal (4, 12, 26, 32). During non-canonical NLRP3 inflammasome activation, human monocytes recognize intracellular LPS, which activates caspases-4 and −5. These caspases are then responsible for caspase-1 activation and the cleavage of IL-1β (2, 4, 33). Human monocytes can also activate an alternative inflammasome, in which extracellular LPS stimulation alone signals through caspase-8, which contributes to NLRP3 inflammasome activation through caspase-8 interactions with FADD and RIPK1 (12). The alternative NLRP3 inflammasome also induces IL-1β release independent of cell death or potassium efflux, two markers of the canonical NLRP3 inflammasome (12).

To further investigate the requirements for inflammasome activation during infection of human cells, we infected primary human peripheral blood monocytes with the protozoan parasite *Toxoplasma gondii* (*T. gondii*), which induces the release of IL-1β (34). *T. gondii* is a member of the phylum *Apicomplexa* and closely related to *Plasmodium* species. It is a eukaryotic single-celled obligate intracellular parasite that is estimated to infect one-third of the global population (35). *T. gondii* infection is life-threating for immunocompromised individuals and developing fetuses, and can also cause morbidity in immunocompetent hosts, as it is a leading cause of hospitalization due to a foodborne pathogen in the United States (36–38). CD4^+^ and CD8^+^ T cells are required for protection against *T. gondii* infection through the production of IFN-γ, but innate immune cells, such as monocytes also significantly contribute to immune control of infection (39–41). Monocytes are among the first cells recruited to sites of infection. They are also preferentially infected by the parasite compared to other PBMCs and once infected, rely on the NLRP3 inflammasome to produce and release bioactive IL-1β (34, 42, 43).

We previously demonstrated that IL-1β is released from *T. gondii*-infected human monocytes in a manner dependent on priming through a Syk-PKCδ-CARD9-MALT1-NF-κB signaling pathway, and activation of an NLRP3 inflammasome that requires caspase-1, ASC, and potassium efflux (42, 43). IL-1β was released from these cells via a process that was independent of cell death or GSDMD (42). However, it is still unclear if caspases other than caspase-1 contribute to IL-1β release from *T. gondii*-infected human monocytes, and the mechanism of IL-1β release from *T. gondii*-infected monocytes has yet to be defined. Here we demonstrate that caspase-8, but not caspases-4 or −5, is also required for the IL-1β response from *T. gondii-infected* human monocytes. Caspase-8 was not involved in the synthesis or cleavage of IL-1β, but rather, caspase-8 functioned in a novel mechanism controlling the release of IL-1β from viable cells through a process that was independent of caspase-3 or GSDMD. Interestingly, caspase-8 also contributed to the release of IL-1β from monocytes infected with the closely related apicomplexan parasite, *Neospora caninum*. The studies presented here expand on the inflammatory functions of caspase-8 and define a novel role for caspase-8 in IL-1β release from viable human immune cells infected with protozoan parasites.

## Materials and Methods

### Ethics Statement

Human whole blood was collected by the Institute for Clinical and Translational Science (ICTS) at the University of California, Irvine from healthy adult donors who provided written informed consent. Blood was collected according to the guidelines of and with approval from the University of California, Irvine Institutional Review Board (HS #2017–3753).

### Parasite and Mammalian Cell Culture

PBMCs were isolated from human whole blood by density gradient centrifugation using lymphocyte separation media (MP Biomedicals, Santa Ana, CA). Monocytes were enriched from PBMCs by counterflow elutriation, as previously described (44), and stained for purity after isolation. This protocol typically resulted in >90% pure monocyte cultures (ranging from 85–95%) based on CD11b^+^ and CD3^-^CD20^-^ CD56^-^ staining. Freshly isolated monocytes were resuspended in RPMI 1640 (HyClone, Logan, UT) supplemented with 2 mM L-glutamine, 100 U/ml penicillin, 100 μg/ml streptomycin, and either 10% heat-inactivated FBS (Omega Scientific, Tarzana, CA) (R-10%) or no serum (R-0%). Monocytes were used immediately after isolation for experiments.

The human monocytic THP-1 cell line was cultured in R-10% supplemented with 2 mM L-glutamine, 100 U/ml penicillin, and 100 μg/ml streptomycin. The caspase-1, −4, −5, and −8 KO THP-1 cells were cultured in the same medium supplemented with 2 μg/ml puromycin.

Human foreskin fibroblasts (HFFs; from the lab of Dr. John Boothroyd, Stanford University School of Medicine) were cultured in D-10% medium: DMEM (HyClone) supplemented with 10% heat-inactivated FBS, 2 mM L-glutamine, 100 U/ml penicillin, and 100 μg/ml streptomycin. *T. gondii (Prugniaud)* and *N. caninum* (NC1) tachyzoites were maintained by serial passage in confluent monolayers of HFFs. *T. gondii* constitutively expressing GFP were used (45).

All mammalian and parasite lines were cultured at 37°C in 5% CO_2_ incubators. All cultures were tested bimonthly and confirmed to be free of *Mycoplasma* contamination

### Generation of Knockout (KO) Cell Lines

Empty vector and knockout THP-1 cells were generated using the Lenti-CRISPR-Cas9 system (42). Guide RNAs (sgRNA) targeting human caspase-1, −4, −5, and −8 were cloned into the LentiCRISPR v2 plasmid (Feng Zhang, Addgene plasmid #52961). The sequences for the guide RNAs were as follows: Caspase-1: TAATGAGAGCAAGACGTGTG, Caspase-4: GAGAAACAACCGCACACGCC, Caspase-5: TGGGGCTCACTATGACATCG, Caspase-8: GCCTGGACTACATTCCGCAA. Virus was generated by transfecting the sgRNA plasmid constructs into HEK 293T cells with the psPAX2 packaging (Didier Trono, Addgene plasmid #12260) and pCMV-VSVG envelope (Bob Weinberg, Addgene plasmid #8454) plasmids. Viral supernatants collected at 48 hr post-transfection were used to infect THP-1 cells by spinfection at 1800 rpm for 1 hr. Single-cell knockout clones were generated by limiting dilution in 96-well plates under puromycin selection. Single-cell clones were sequenced after PCR amplification of a 500 bp region near the Cas9 binding site. Interference of CRISPR edits (ICE) analysis software (Synthego) was used to characterize the indel for each clone (46). A caspase-4 clone containing a 1 base pair (bp) deletion at the CARD binding domain, a caspase-5 clone containing a 20 bp deletion in the p20 subunit, a caspase-8 clone containing a 5 bp deletion in the death effector domain, and a caspase-1 clone containing a 1 bp insertion in the substrate binding pocket were used in subsequent experiments. All the caspase KO clones were screened via Western blot for the presence of Cas9 and the absence of the gene targeted for deletion. Since we could not detect caspase-5 by Western blot using several commercially available antibodies, as has been previously reported (47), the KO clone was verified by sequencing.

### Infection and Stimulation of Monocytes

Primary human monocytes and monocytic THP-1 cells were resuspended in R-0% or R-10% medium directly after isolation and incubated with small molecule inhibitors or equal volumes of the vehicle control for 40 minutes at 37°C. *T. gondii*-infected HFFs were washed with D-10% medium, scraped, and syringe lysed. Lysed tachyzoites were washed with medium, passed through a 5-μm filter (EMD Millipore, Billerica, MA), and washed with medium again. This resulted in parasite cultures that were free of host cell debris and soluble factors. Purified *T. gondii* tachyzoites were immediately added to host cells at a multiplicity of infection (MOI) of 2. All infections were performed with a GFP-expressing type II *Prugniaud* strain of *T. gondii*. “Mock” infections were samples in which an equivalent volume of culture medium without parasites was added to the cells. Experiments with primary human monocytes were harvested after 4 hours post-infection (hpi) and those with THP-1 cells were harvested at 16-18 hpi.

Cells were stimulated with 100 ng/ml ultrapure *E. coli* LPS (List Biological Laboratories, Campbell, CA) and then 5 mM ATP (Sigma-Aldrich, St. Louis, MO) for the last 30 min of culture, as indicated. “Mock” treatment was the addition of the equivalent volume of media (without parasites or LPS) to cells. At the indicated time point, monocytes were pelleted by centrifugation at 500 x g for 5 min. Collected cells were stained, fixed, or lysed accordingly, as described below.

### Inhibitors

MCC950 (Adipogen, San Diego, CA) was resuspended in deionized water. IETD (R&D Systems, Minneapolis, MN), Ac-Tyr-Val-Ala-Asp-chloromethylketone (Ac-YVAD-CMK or YVAD) (Cayman Chemical, Ann Arbor, MI), ZVAD (Selleckchem, Houston, TX) and Z-DEVD-FMK (Torcis Biosciences, Bristol, UK) were all resuspended in DMSO. Monocytes were treated with the inhibitors or with an equivalent volume of the appropriate vehicle, for 30 min at 37°C and then infected or stimulated as described above. Nec1 (Cayman Chemical, Ann Arbor, MI) or an equal volume of DMSO for the vehicle control was added to cells 30 minutes before YVAD or IETD treatment. MCC950, IETD, Nec1, YVAD, and ZVAD were all used at concentrations that did not induce cell death or reduce infection efficiency.

### Flow Cytometry

To measure the efficiency of *T. gondii* infections, cells were harvested at the indicated time points and immediately analyzed by flow cytometry to determine the percent of GFP^+^ (*T. gondii*-infected) cells. To measure cell viability, cells were harvested, washed and resuspended in FACS buffer (2% FBS in PBS) containing propidium iodide (eBioscience, San Diego, CA) and analyzed by flow cytometry without fixation.

Samples were analyzed on a BD FACSCalibur or Agilent Novocyte 3000 flow cytometer. Data were analyzed using FlowJo software (TreeStar, Ashland, OR). Cells were first identified based on their forward and side scatter profile and subsequently analyzed for cell surface marker expression, intracellular cytokine expression, or GFP signal.

### Quantitative Real-Time PCR (qPCR)

At the harvest time point, total RNA was harvested using the RNeasy Kit (QIAGEN, Germantown, MD) and treated with DNase I (Life Technologies, Carlsbad, CA) to remove any contaminating genomic DNA. cDNA was synthesized using the Superscript III First-Strand Synthesis kit (Life Technologies), according to the manufacturer’s instructions, and subsequently used as template in quantitative real-time PCR (qPCR). qPCR was performed in triplicate using a Bio-Rad iCycler PCR system (Bio-Rad, Hercules, CA) and iTaq Universal SYBR Green Supermix (Bio-Rad). Previously published sequences for *pro-IL-1ϐ* (48) and *GAPDH* (49) primers were used. All primer pairs spanned intron-exon boundaries whenever possible and bound to all isoforms of the gene, where applicable. All primers were commercially synthesized by Integrated DNA Technologies (Coralville, IA). qPCR data were analyzed using the threshold cycle method, as previously described (50), and gene expression data are shown normalized to that of the housekeeping gene *GAPDH*. In all qPCR assays, cDNA generated in the absence of reverse transcriptase, as well as water in the place of DNA template, were used as negative controls, and these samples were confirmed to have no amplification.

### ELISA

Concentrations of human IL-1β, TNF-α and IL-6 protein were quantified using ELISA MAX Deluxe kits (BioLegend, San Diego, CA), according to the manufacturer’s instructions. Cytokines in the supernatant were diluted in blocking buffer and added directly to plates. Intracellular IL-1β was measured by lysing cells through 3 quick freeze-thaw cycles using liquid nitrogen and a 37°C water bath followed by manual disruption of the cell pellet through pipetting. The resulting lysate was centrifuged for 10 minutes at 14,000 x g to remove cellular debris, and the lysate was diluted in blocking buffer before addition to the plate. Signal from ELISA plates was read with a Spectra Max Plus 384 plate reader (Molecular Devices, San Jose, CA) using SoftMax Pro Version 5 software (molecular Devices), and the threshold of detection was 0.5 pg/ml.

### Caspase Activity Assay

Caspase-1 activity was quantified using a Caspase-Glo 1 Inflammasome Assay kit (Promega, Madison, WI). THP-1 cells were treated and harvested as described above. 50,000 cells resuspended in 100 μL of media or 100 μL of supernatant were added in triplicate to opaque 96-well plates. 100 μL of caspase-1 activity detection reagent was added to 2 of the 3 replicates for each sample while 100 μL of the caspase-1 activity reagent containing the caspase-1 inhibitor YVAD was added to the 3^rd^ replicate. The plate was sealed, mixed by shaking for 2 minutes and left to incubate at room temperature in the dark for 1 hour. Luminescence in the plate was then read with a SpectraMax i3x (Molecular Devices, San Jose, CA) and analyzed using SoftMax Pro Version 5 software (Molecular Devices, San Jose, CA). Caspase-1 activity was quantified by subtracting the luminescence in the wells containing detection reagent plus YVAD from the average luminescence of the wells containing only the caspase-1 activity reagent.

### Western Blots

At the harvest time point, cells were lysed by addition of 2X Laemmli buffer containing 10% 2-ME or a TNE lysis buffer (50 mM Tris-HCl pH 8.0, 150 mM NaCl, 1% NP-40, 3 mM EDTA). Each Buffer also contained a protease inhibitor cocktail (1 mM Na_3_VO_4_ pH 8.0, 5 mM NaF, 10 mM Na_4_PO_7_). For experiments in which supernatant was analyzed, serum-free R-0% medium was used during the infection, and supernatant was concentrated using Amicon Ultra Centrifugal filters (EMD Millipore, Burlington, MA), according to the manufacturer’s instructions. Concentrated supernatants were diluted with 2X Laemmli buffer containing 10% 2-ME. Samples were boiled at 100°C for 10 to 15 min, and then centrifuged for 10 minutes at 14,000 x g to remove cellular debris. Samples were separated by SDS-PAGE and transferred to polyvinylidene difluoride (PVDF) membranes (Bio-Rad, Hercules, CA) for immunoblotting. Membranes were blocked for 1 h at room temperature (RT) with blocking buffer: 5% non-fat milk or 5% bovine serum albumin (BSA) (Fisher Bioreagents). Membranes were then incubated with primary antibodies diluted in blocking buffer for 3 h at RT or overnight at 4°C. Membranes were probed with antibodies against caspase-1 (ab179515; Abcam), caspase-4 (#4450; Cell Signaling Technology), caspase-8 (#9746; Cell Signaling Technology), or β-actin (AC-15; Sigma-Aldrich). Membranes were blotted for pro-IL-1β and IL-1β (3ZD from the National Cancer Institute Biological Resources Branch) using the SNAP i.d. Protein Detection System (EMD Millipore), according to the manufacturer’s instructions. Primary Abs were followed by HRP-conjugated secondary Abs (BioLegend), and membranes were developed with SuperSignal West Femto Maximum Sensitivity Substrate (Thermo Fisher Scientific, Carlsbad, CA), Amersham ECL Prime Western Blotting Detection Reagent (GE Healthcare, Little Chalfont, U.K.) substrate, Clarity Western ECL Substrate (Bio-Rad, Hercules, CA), ECL Prime Substrate (Thermo Fisher Scientific, Carlsbad, CA) or Clarity Western (Bio-Rad, Hercules, CA). Signal was detected using a Nikon camera, as previously described (51) or with x-ray film development with Amersham Hyperfilm ECL (Global Life Sciences, London, UK). Quantification analysis of blots was performed using ImageJ software.

### Statistics

Statistical analyses were performed using GraphPad Instat software. Analysis of variance (ANOVA) followed by Tukey’s or Bonferroni’s test, as indicated, were used for comparison between means. Differences were considered significant when the P value was <0.05.

## Results

### Caspases-1 and −8, but not caspases-4 or −5, contribute to IL-1β release from *T. gondii*-infected human monocytes

Caspases-1, −4, −5 and −8 have each been shown to contribute to IL-1β release from human monocytes and macrophages during activation of the canonical, noncanonical or alternative NLRP3 inflammasomes during LPS stimulation or pathogen infection (4, 10, 12, 25, 52). We and others have previously reported a role for caspase-1 in *T. gondii-induced* IL-1β production (34, 42, 43, 53). To examine a role for other caspases, caspase-1, −4, −5, and −8 knockout (KO) cells were generated using CRISPR/Cas9 genome editing in the THP-1 human monocytic cell line. Control empty vector (EV) THP-1 cells lacking the targeting guide RNAs were also generated for comparison. After the selection and clonal expansion of the stable KO and EV cell lines, the clones were sequenced to identify the mutations introduced by the CRISPR/Cas9 system (Supp. Fig. 1A). Western blot analysis of the EV and KO clones demonstrated the absence of the respective caspases and the presence of Cas9 in the KO cells (Supp. Fig. 1B).

Clonal lines of the caspase KO and EV cells were infected with a type II (*Prugniaud*) GFP-expressing strain of *T. gondii* or exposed to an equal volume of media (mock), and the supernatants were analyzed for IL-1ß levels by ELISA at 18 hours post-infection (hpi). *T. gondii*-infected caspase-1 and −8 KO cells, but not caspase-4 or −5 KO cells, released significantly less IL-1β compared to the infected EV cells (Fig. 1A). The reduced IL-1β release was not due to reduced infection of the KO lines, as there was no difference in the percent of infected (GFP^+^) cells in each condition (Supp. Fig. 1C). Since caspases can contribute to cell death pathways, we evaluated the effect of infection on cell death in the EV and caspase-1 and −8 KO cells by staining with propidium iodide (PI). There were no significant differences in viability among the infected and mock-treated conditions in either the EV or caspase KO lines (Fig. 1B). As a positive control for PI staining, EV cells were stimulated with LPS and ATP, which activates the canonical NLRP3 inflammasome and induces pyroptosis, and a significant increase in PI^+^ cells was observed (Fig. 1B). As an additional control to evaluate the caspase-8 KO cells in response to well-characterized stimuli, we also treated them with LPS (Fig. 1C) or LPS and ATP (Fig. 1D). As expected, the caspase-8 KO cells released less IL-1β than the EV cells during LPS stimulation (Fig. 1C), but caspase-8 was dispensable for IL-1β release during LPS and ATP stimulation (Fig. 1D), reinforcing that caspase-8 is not required for IL-1β release during activation of the canonical NLRP3 inflammasome (12). Finally, the role of caspase-8 in cytokine release appeared to be specific to IL-1β, as caspase-8 KO cells and EV cells released comparable levels of other proinflammatory cytokines, including IL-6 and TNF-α, during *T. gondii* infection (Fig. 1E).

**Fig 1.**
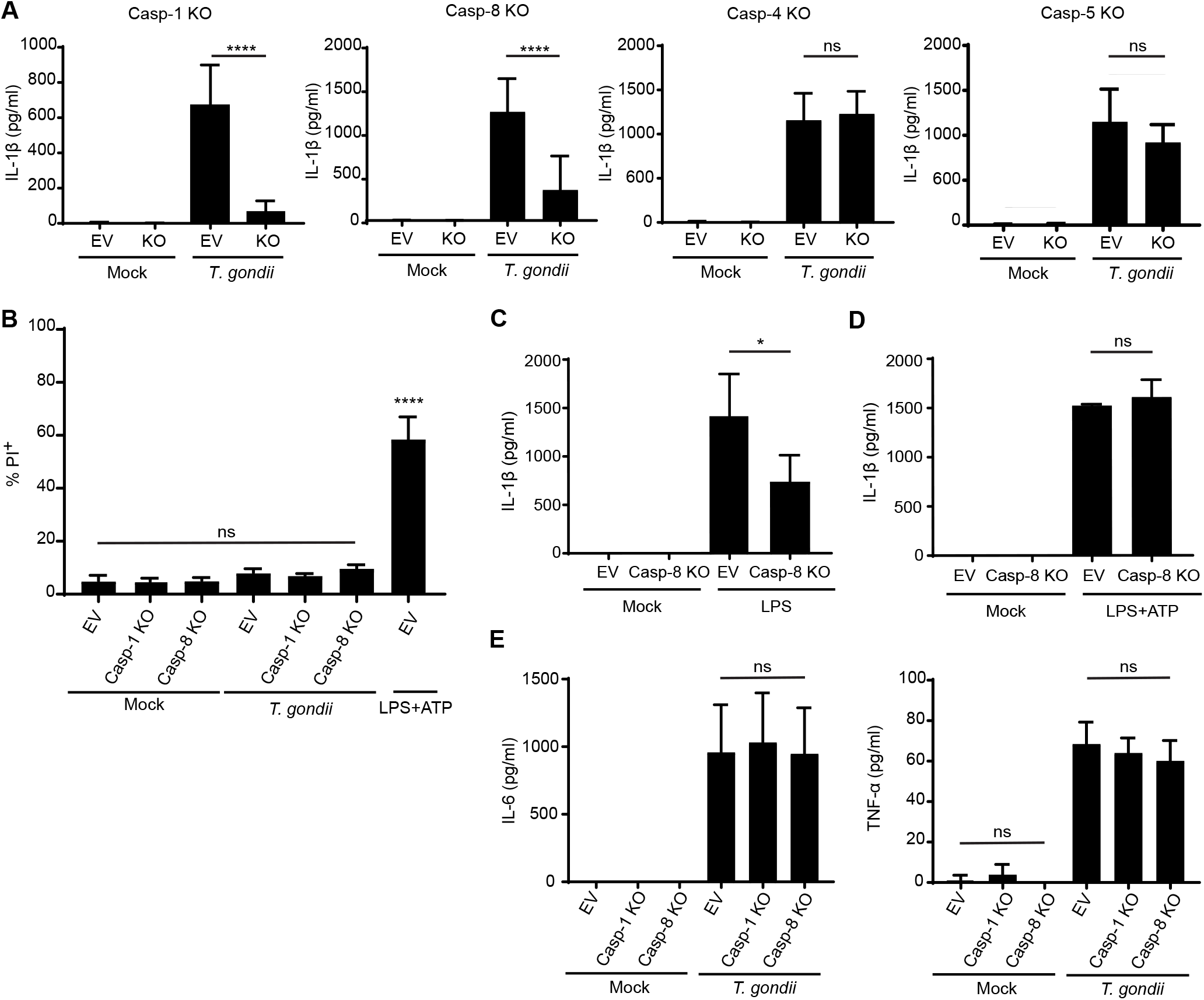
IL-1β release during *T. gondii* infection of THP-1 cells is dependent on caspase-1 and caspase-8. EV and caspase KO THP-1 cells were mock treated or infected with *T. gondii* at a MOI of 2. **(A)** IL-1β in the supernatant was detected by ELISA. n = 4-8 experiments for each set of caspase KO cells. **(B)** EV and caspase KO THP-1 cells were mock treated, infected with *T. gondii*, or stimulated with LPS and ATP. Cells were stained with propidium iodide (PI) and analyzed by flow cytometry for the percent of PI^+^ cells. n = 7 experiments. **(C, D)** EV and caspase-8 KO cells were mock treated or stimulated with (C) LPS alone or (D) LPS and ATP as above. IL-1β in the supernatant was detected by ELISA. n = 3-4 experiments per condition. **(E)** EV and caspase-8 KO cells were mock treated or infected with *T. gondii*. TNF-α and IL-6 in the supernatant were detected by ELISA. n = 4 experiments for each cytokine. Values are expressed as the mean ± SD, ns (not significant), * *P* < 0.05, **** *P* < 0.0001 (one-way ANOVA followed by a Tukey post-test).

### Primary human monocytes infected with *T. gondii* release IL-1β in a caspase-8-dependent manner

Given the reduction in IL-1β release observed during *T. gondii* infection of caspase-8 KO THP-1 cells, we investigated whether primary human monocytes also relied on caspase-8 for IL-1β release during infection. These cells were isolated from healthy blood donors as previously described (42, 43). Since the short lifespan of primary monocytes limits the genetic tools available to manipulate their activity, the caspase-8-specific inhibitor IETD was used. In addition to its role in apoptosis, caspase-8 also functions in preventing necroptosis by inhibiting RIPK1 (8). Therefore, caspase-8-inhibited cells were also treated with Necrostatin-1 (Nec1), a RIPK1 inhibitor, to prevent necroptosis in the absence of caspase-8 activity. As we have previously reported, *T. gondii* infection of primary human monocytes from multiple independent blood donors induced the release of IL-1β by 4 hpi (Fig. 2A), a more rapid timescale than in THP-1 cells (34, 42, 43). Nec1 treatment had no effect on IL-1β release in mock or infected conditions. In contrast, *T. gondii*-infected cells treated with IETD and Nec1 released significantly less IL-1β compared to cells treated with Nec1 or vehicle control alone (Fig. 2A). Importantly, there was no effect of IETD and Nec1 inhibition on the parasite’s ability to infect these cells, compared to DMSO, IETD, or Nec1 treatment alone (Supp Fig. 2A). To detect the release of bioactive IL-1β from cells, Western blot was used to probe for mature IL-1β (17 kD) in the cell supernatants. The combination of caspase-8 and necroptosis inhibition (IETD + Nec1) significantly reduced mature IL-1β in the supernatant (Fig. 2B), corroborating the ELISA results (Fig. 2A). As a positive control, primary human monocytes were treated with LPS, which induces IL-1β release via a caspase-8-dependent mechanism (12). As expected, Nec1 and IETD treatment also reduced IL-1β release during LPS stimulation of primary monocytes (Fig. 2C). Inhibiting caspase-1 with YVAD or caspase-8 with IETD + Nec1 resulted in similar reductions in IL-1β release (Supp. Fig. 2B), and dual inhibition of caspase-1 and caspase-8 further reduced IL-1β release (Supp. Fig. 2B). These data suggest that caspase-1 and caspase-8 may function in non-redundant pathways.

**Figure 2.**
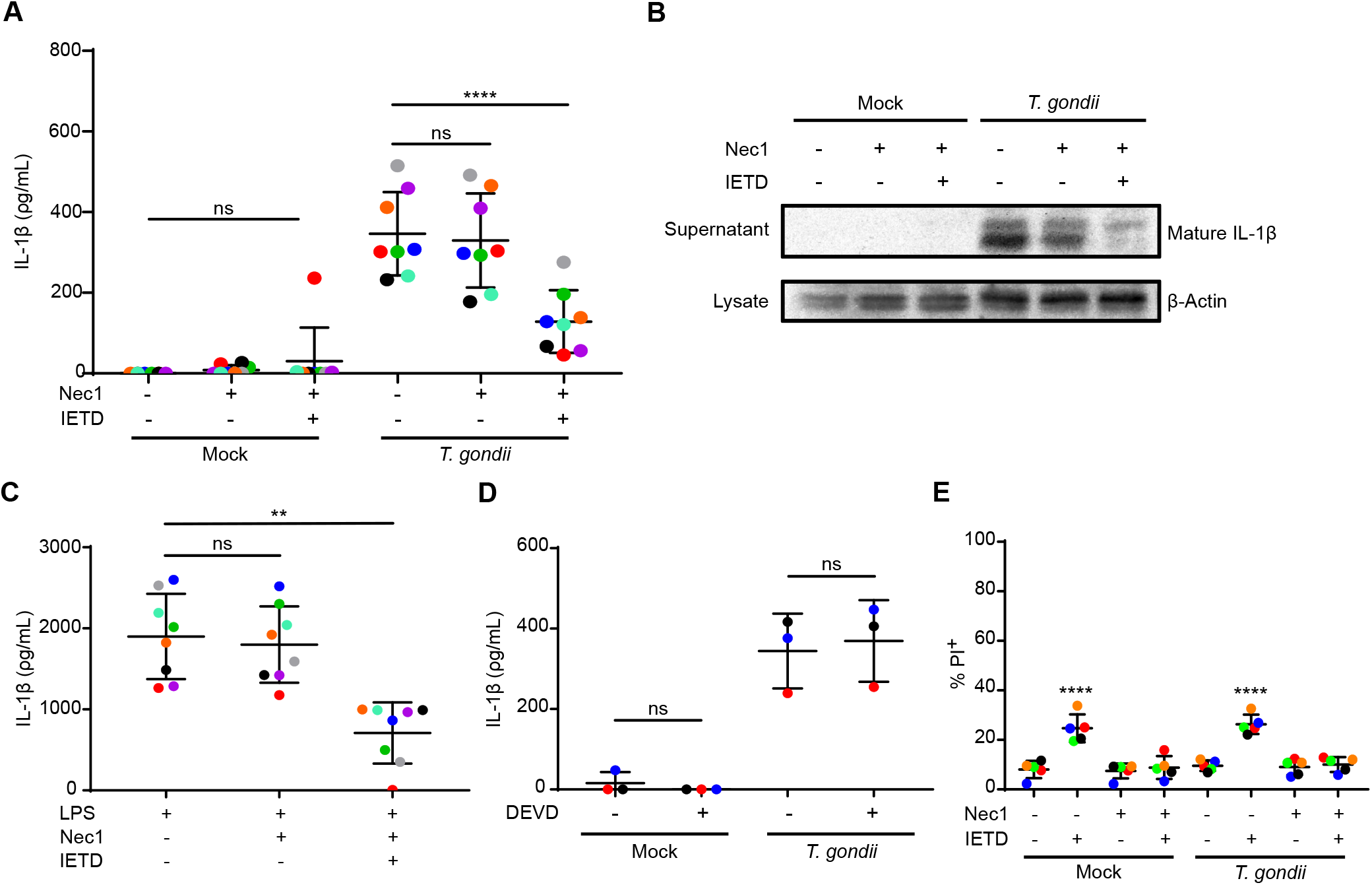
Human peripheral blood monocytes utilize caspase-8 for IL-1β release during *T. gondii* infection. **(A, B)** Primary human monocytes were pretreated with a vehicle control, Nec1, or Nec1 and IETD and then mock treated or infected with *T. gondii* at a MOI of 2. **(A)** IL-1β in the supernatant was quantified by ELISA. Each colored dot represents a separate human blood donor. n = 9 experiments with independent blood donors. **(B)** Mature IL-1β in supernatants and β-actin from corresponding cell lysates were detected by Western blot. **(C)** Primary human monocytes were pretreated with a vehicle control, Nec1, or Nec1 and IETD and stimulated with LPS. Supernatants were used for detection of IL-1β by ELISA. n = 8 experiments. **(D)** Primary human monocytes were pre-treated with DEVD or vehicle and then infected with *T. gondii* or mock infected. IL-1β was detected in supernatants by ELISA. n = 3 experiments. **(E)** Primary human monocytes were treated as described above, the cells were stained with PI, and the percentage of PI^+^ cells was quantified by flow cytometry. n = 5 experiments. Values are expressed as the mean ± SD, ns (not significant), * *P* < 0.05, ** *P* < 0.01, **** *P* < 0.0001 (one-way ANOVA followed by a Tukey post-test).

Caspase-8 has been implicated in IL-1β release through the activation of caspase-3, which subsequently cleaves gasdermin E and induces cell death and IL-1β release (10). However, pretreatment of monocytes with the caspase-3 inhibitor DEVD did not reduce IL-1β release during *T. gondii* infection (Fig. 2D), suggesting that caspase-3 is not involved in *T. gondii*-induced IL-1β release. Moreover, to evaluate the potential effect of Nec1 and IETD treatment on monocyte viability during *T. gondii* infection, cell death was analyzed by the uptake of PI. IETD treatment of mock or infected cells in the absence of Nec1 inhibition resulted in elevated cell death, as expected, since caspase-8 also functions in inhibiting RIPK1/3-mediated necroptosis. The addition of Nec-1 to IETD-treated cells restored their viability (Fig. 2E). *T. gondii* infection did not result in increased cell death, as we have previously reported (42). Collectively, these data indicate that the reduction in IL-1β release in caspase-8-inhibited primary cells was not due to an effect on cell death.

Likewise, infection of murine macrophages with another closely related apicomplexan parasite, *Neospora caninum* (*N. caninum*), has also been shown to induce IL-1β release through activation of the NLRP3 inflammasome (54, 55). We next investigated a potential role for caspase-8 in IL-1β release during *N. caninum* infection. Treatment of primary human monocytes with inhibitors of NLRP3 (MCC950), caspase-1 (YVAD), caspase-8 (IETD+Nec1), or with a pan-caspase inhibitor (ZVAD) prior to infection with *N. caninum* also reduced IL-1β release, indicating that this release was also dependent on NLRP3, and caspases-1 and −8 (Supp. Fig. 3). These data suggest that caspase-8 is required for IL-1β release from primary human monocytes during infection with multiple apicomplexan parasites.

### Caspase-8 is not required for pro-IL-1β synthesis during *T. gondii* infection of human monocytes

Although caspase-8 inhibition or deficiency reduced the detection of mature IL-1β in the supernatants of both primary human monocytes and THP-1 cells during *T. gondii* infection, the mechanism by which this occurred was unclear. We considered the possibility of caspase-8 functioning either in inducing IL-1β synthesis (9, 56), or participating in the cleavage of IL-1β via activation of the NLRP3 inflammasome (11, 13, 57). Since caspase-8 contributes to *IL-1ϐ* transcript production in murine bone marrow-derived macrophages (BMDMs) during *T. gondii* infection (9, 56), we first examined whether caspase-8 performs a similar function in human monocytes. As expected, *IL-1ϐ* transcripts were induced in THP-1 cells during *T. gondii* infection; however, there was no significant difference in *IL-1ϐ* mRNA in caspase-8 KO cells in response to *T. gondii* (Fig. 3A). In addition, pro-IL-1β protein levels in cell lysates of caspase-8 KO THP-1 cells (Fig. 3B) or caspase-8-inhibited primary monocytes (Fig. 3C) were not significantly different from those in infected EV THP-1 cells or mock-treated primary monocytes. These data strongly suggest that caspase-8 does not contribute to the synthesis of *IL-1ϐ* transcripts or the production of pro-IL-1β protein during *T. gondii* infection of human monocytes.

**Figure 3.**
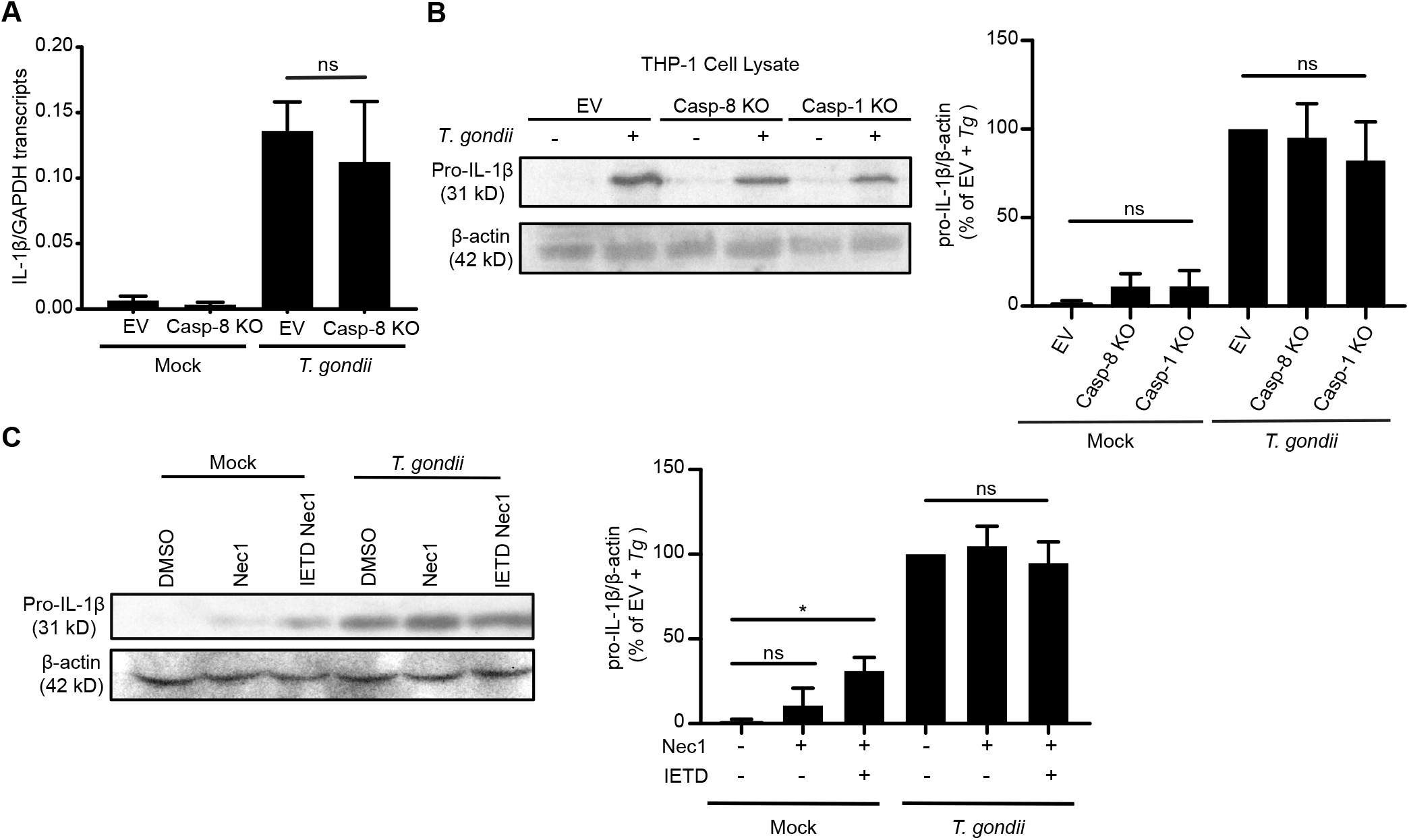
Caspase-8 is not required for pro-IL-1β synthesis during *T. gondii* infection of human monocytes. **(A, B)** EV and caspase-8 KO cells were mock treated or infected with *T. gondii* at a MOI of 2. **(A)** *IL-1ϐ* and *GAPDH* transcript levels were measured by qPCR at 6 hpi. A representative experiment (of n = 3) is shown. **(B)** EV, caspase-1, and caspase-8 KO cells were harvested, and cell lysates were probed for pro-IL-1β by Western blot. A representative blot is shown (left), and pro-IL-1β levels were measured by densitometry (right) from n = 4 experiments. **(C)** Primary human monocytes were treated with vehicle, Nec1, or Nec1 and IETD, and cell lysates were examined for pro-IL-1β by Western blot. A representative blot of is shown (left), and pro-IL-1β levels were measured by densitometry (right) from n = 3 experiments. Values are expressed as the mean ± SD, ns (not significant), * *P* < 0.05 (one-way ANOVA followed by a Tukey post-test).

### Caspase-8 is not required for caspase-1 activity or IL-1β cleavage during *T. gondii* infection of human monocytes

Caspase-8 can function as an inflammatory caspase by activating the non-canonical or alternative NLRP3 inflammasome to induce caspase-1 activation, or, in the absence of caspase-1 activity, caspase-8 can directly cleave IL-1β and gasdermin D (10–13, 58). To investigate whether caspase-8 in *T. gondii*-infected human monocytes activates the inflammasome, a luminescence-based caspase activity assay was used to probe for caspase-1 activity in the lysates of *T. gondii*-infected EV control cells and caspase-1 and caspase-8 KO cells. An increase in caspase-1 activity was detected during both *T. gondii* infection and during LPS and ATP dual stimulation of EV control cells, but not in caspase-1 KO cells, as expected (Fig. 4A). Notably, there were similar levels of active caspase-1 in caspase-8 KO cells compared to EV cells during infection (Fig. 4A). These data indicate that caspase-1 activation can occur in the absence of caspase-8 during *T. gondii* infection and suggest that NLRP3 inflammasome activation is intact in caspase-8 KO cells.

**Figure 4.**
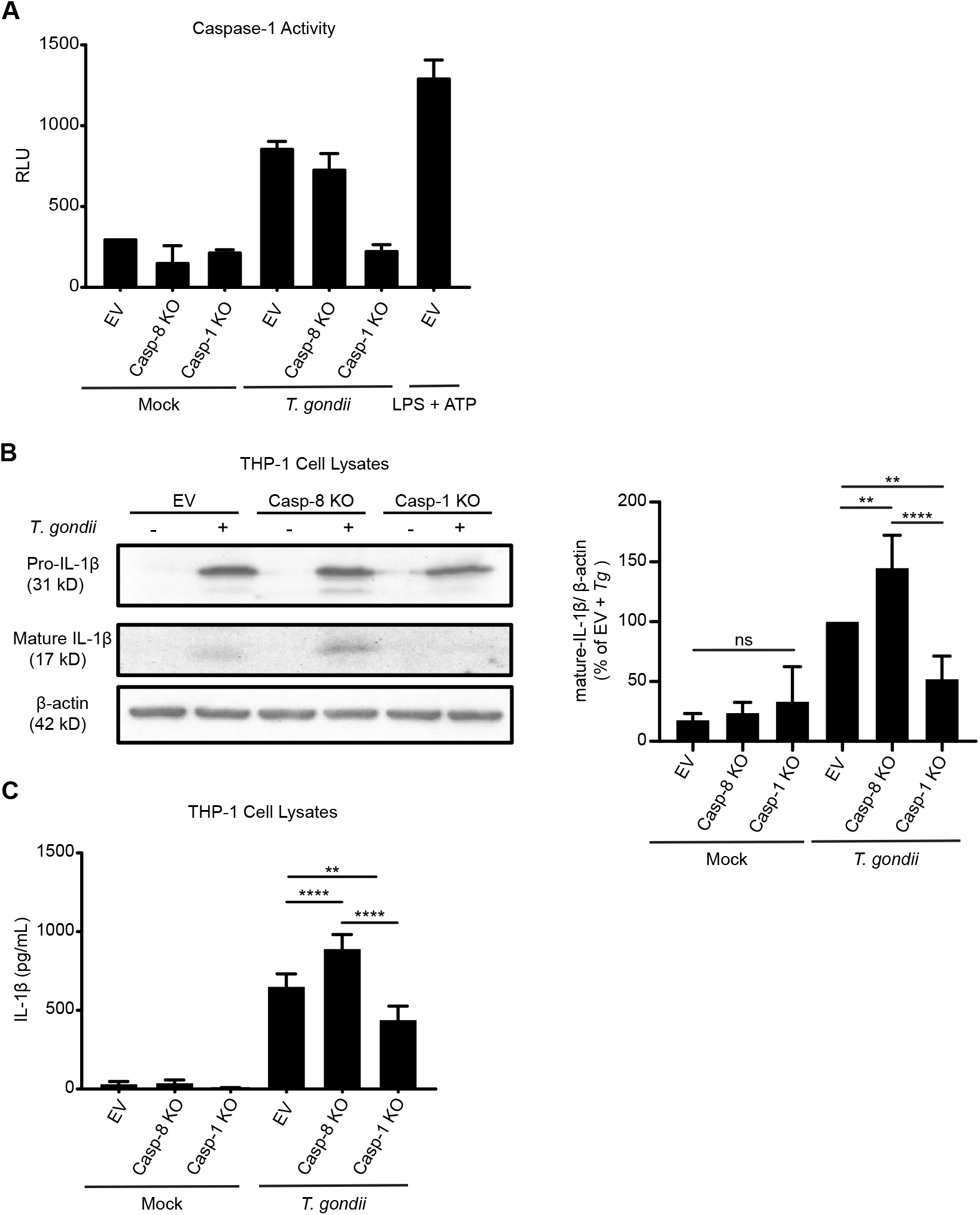
Caspase-8 is not required for caspase-1 activity or IL-1β cleavage during *T. gondii* infection. EV, caspase-1, and caspase-8 KO cells were mock treated, stimulated with LPS and ATP, or infected with *T. gondii* at a MOI of 2. **(A)** Caspase-1 activity form cell lysates was quantified. A representative experiment of n = 2 is shown. **(B)** IL-1β and β-actin in cell lysates were probed by Western blot. A representative blot is shown (left) and mature-IL-1β levels from n = 3 blots were measured by densitometry. **(C)** Cell lysates were probed for IL-1β by ELISA. n = 4 experiments. Values are expressed as the mean ± SD, ns (not significant), ** *P* < 0.01, *** *P* < 0.001, **** *P* < 0.0001 (one-way ANOVA followed by a Tukey post-test).

To directly test whether caspase-8 is required for IL-1β cleavage, the EV, caspase-8, and caspase-1 KO cells were mock treated or infected with *T. gondii*, and the cell lysates were examined for pro- and mature IL-1β by Western blot. Levels of pro-IL-1β were equivalent between the infected caspase-8 KO and EV THP-1 cells. Interestingly, there was no reduction in the cleavage of IL-1β in the infected caspase-8 KO cells, but rather, a significant accumulation of cleaved IL-1β in the caspase-8 KO cells was detected compared to infected EV THP-1 cells (Fig. 4B). These results were corroborated by ELISAs measuring IL-1β levels inside infected EV and caspase-8 KO cells: lysates from infected caspase-8 KO cells contained significantly more IL-1β than infected EV control cells (Fig. 4C). Considering that *T. gondii*-infected human monocytes appeared to accumulate mature IL-1β intracellularly, these data strongly suggest that caspase-8 is not essential for IL-1β synthesis or cleavage in *T. gondii*-infected human monocytes, but instead likely contributes to the *release* of mature IL-1β from viable monocytes.

### Caspase-8 influences the release of mature IL-1β and active caspase-1 from *T. gondii*-infected cells

To further investigate the importance of caspase-8 in the release of IL-1β from cells, the supernatants of *T. gondii*-infected caspase-1 and caspase-8 KO and EV THP-1 cells were examined for mature IL-1β by Western blot. As expected, mature IL-1β was not detected in supernatants of infected caspase-1 KO cells (Fig. 5A). In addition, there was a significant decrease in the amount of both pro- and mature IL-1β detected in the supernatants of infected caspase-8 KO cells compared to infected empty vector cells (Fig. 5A). The fact that a low level of mature IL-1β was detectable in the supernatants of caspase-8 KO cells, whereas caspase-1 KO cells had no detectable mature IL-1β in the supernatant (Fig. 5A), indicates that a small amount of mature IL-1β may be released independent of caspase-8.

**Figure 5.**
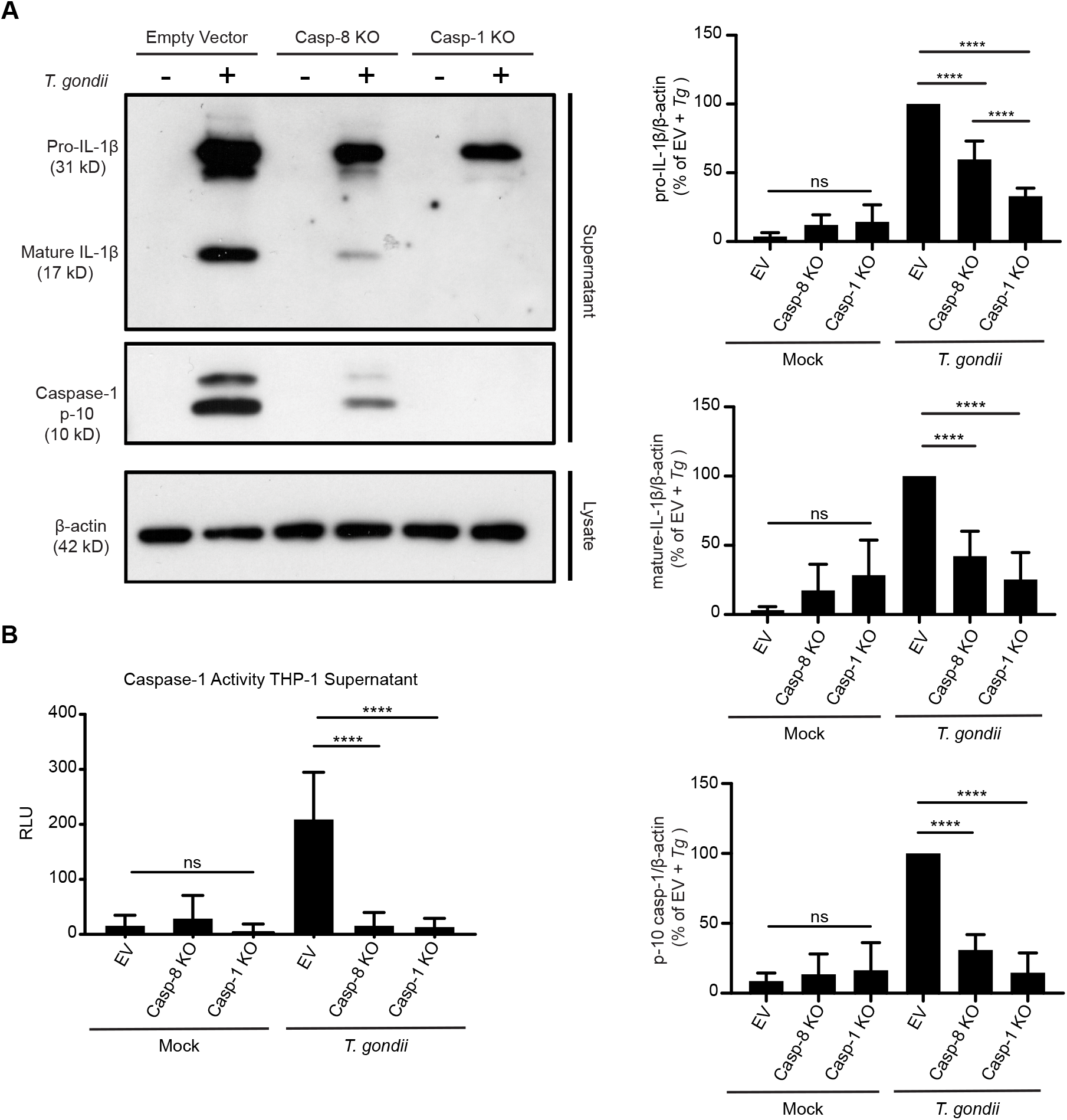
Caspase-8 contributes to the release of mature IL-1β and active caspase-1 from *T. gondii* infected cells. EV, caspase-1 KO, and caspase-8 KO cells were mock treated or infected with *T. gondii* at an MOI of 2. **(A)** Pro- and mature IL-1β and active caspase-1 (p10) in supernatants, and β-actin from corresponding cell lysates were visualized by Western blot. A representative blot is shown (left), and densitometry of n = 5 blots (for IL-1β) and 4 blots (for caspase-1) are quantified. **(B)** Supernatants were evaluated for caspase-1 activity. n = 3 experiments. Values are expressed as the mean ± SD, ns (not significant), **** *P* < 0.0001 (one-way ANOVA followed by a Tukey post-test).

Prior work by our lab and others has demonstrated the release of active caspase-1 into the supernatant of cells upon inflammasome activation. Intriguingly, when supernatants were probed for caspase-1, the cleaved p10 fragment of caspase-1 could be detected in the supernatants of infected EV cells, but significantly less caspase-1 p10 was found in the supernatants of infected caspase-8 KO cells (Fig. 5A, right). We next examined caspase-1 activity in the supernatants of infected EV, caspase-1 KO, and caspase-8 KO cells using a luminescence-based caspase-1 activity assay, and similarly, significantly less active caspase-1 was detected in the supernatants of caspase-8 KO cells (Fig. 5B). Since caspase-8 deficiency did not significantly decrease the levels of active caspase-1 in the cell lysates (Fig. 4A), these data indicate that caspase-8 may affect the release of other inflammasome machinery, in addition to the release of IL-1β itself. Collectively, these data support a role for caspase-8-dependent IL-1β release that is independent of pyroptosis or cell membrane pore formation.

## Discussion

Recent research has revealed that caspase-8 and the NLRP3 inflammasome are at a nexus between the induction of inflammation and the regulation of several types of cell death (7, 8, 59). In addition to its functions in regulating apoptosis and necroptosis, caspase-8 can contribute to both NLRP3 inflammasome priming and activation (9, 11, 56, 60). In instances of caspase-1 inactivation, caspase-8 can also induce IL-1β and GSDMD cleavage, activation of caspases −3 and −7 leading to GSDME cleavage and IL-1 release, and the induction of pyroptosis (10, 58, 60). Here we show caspase-8, but not caspase-4 or −5, is a critical mediator of IL-1β release from *T. gondii*-infected human monocytes through a mechanism that is independent of cell death or GSDMD. We did not observe an effect of caspase-8 inhibition on *IL-1ϐ* transcripts or protein production, induction of NLRP3 inflammasome activation, as measured by caspase-1 activity, or the cleavage of pro-IL-1β to mature IL-1β. Instead, loss of caspase-8 function resulted in reduced IL-1β release, as mature IL-1β accumulated intracellularly in *T. gondii*-infected caspase-8 KO cells.

Since the isolation of IL-1β in 1977 (14) and its cloning in 1985 (61), the mechanism by which this inflammatory cytokine is released from cells remained unresolved for almost 20 years. Eventually IL-1β was found to be released through a caspase-1-mediated inflammatory cell death named pyroptosis (29). There is now mounting evidence that IL-1β can also be released from viable cells through several different mechanisms, including exosome release and secretory autophagy (26, 62–64). Although caspase-8 has not previously been found to contribute to the release of IL-1β, caspase-8 is involved in the secretion of inflammatory lysyl-tRNA synthetase in exosomes (65). It is also possible that caspase-8 may cleave an as-yet uncharacterized substrate involved in the trafficking of IL-1β into exosomes. A growing body of research has shown crosstalk between caspase-8 activity and autophagy related proteins (66–68). Autophagy is a mechanism for bulk turnover of intracellular components in response to cell stress and involves the formation of autophagosomes, double membrane-bound vesicles, that can surround intracellular proteins (69). Caspase-8 localization with autophagosomes has been shown in T cells (66), and IL-1β secretion through autophagosomes has been investigated (64). However, caspase-8 participation in this mechanism of IL-1β release during *T. gondii* infection of human monocytes is still unknown.

There is a growing appreciation that multiple caspases can function in inflammasome activation and IL-1β cleavage. In addition to the canonical NLRP3 inflammasome, caspases-4 and −5 were found to regulate a one-step non-canonical inflammasome mediating IL-1a and IL-1β release in response to LPS stimulation (1–4). An “alternative” NLRP3 inflammasome was described that requires caspases-1 and −8 for cell death-independent release of IL-1β (12). *T. gondii* infection of human monocytes appears to activate the NLRP3 inflammasome, which requires potassium efflux and caspase-1 (42, 43). However, IL-1β release is independent of cell death and gasdermin D, and instead, is largely dependent on caspase-1 and caspase-8. Moreover, our data suggest that during *T. gondii* infection, caspase-8 plays a unique role: it is not required for activation of the inflammasome, but rather, for release of IL-1β and caspase-1 from cells.

To our knowledge, this function for caspase-8 in mediating IL-1β release independent of cell death has not been described previously. The use of human monocytes to probe inflammasome function may have helped to reveal this new role for caspase-8, since human monocytes and neutrophils regulate inflammasome activity differently than murine and human macrophages. Indeed, the non-canonical and alternative NLRP3 inflammasomes were both described in human monocytes (12). In addition, *T. gondii* may activate distinct signaling pathways and mechanisms of inflammasome activity than bacterial or viral pathogens. *T. gondii-derived* factors or parasite-secreted effector proteins may inhibit gasdermin D or other inflammasome-related proteins, thereby allowing infected cells to maintain viability during IL-1β release from the cells.

Together, these findings provide a foothold for initiating investigations into novel functions of caspase-8 and contribute to our understanding of how human monocytes respond to infection with a parasite that causes a large disease burden and morbidity globally.

## Acknowledgements

We would like to thank members of the Lodoen, Andrade, and Morrissette labs, as well as Dr. Eric Pearlman and Dr. David Fruman for helpful discussion on the project. Dr. Andrea Tenner and her lab were also a wonderful help and provided access to their flow cytometer and elutriator. Finally, we are grateful to the administration, nurses and blood donors of the UC Irvine Institute of Clinical and Translational Sciences (ICTS) for maintaining the Healthy Blood Donor program, without which these projects would not be possible.

## Funding

This work was supported by NIH R01AI120846 (to M.B.L.), NIH R21AI156452 (to M.B.L), and NIH T32AI060573-12 (to W.J.P.)

---

**Fig. S1.**
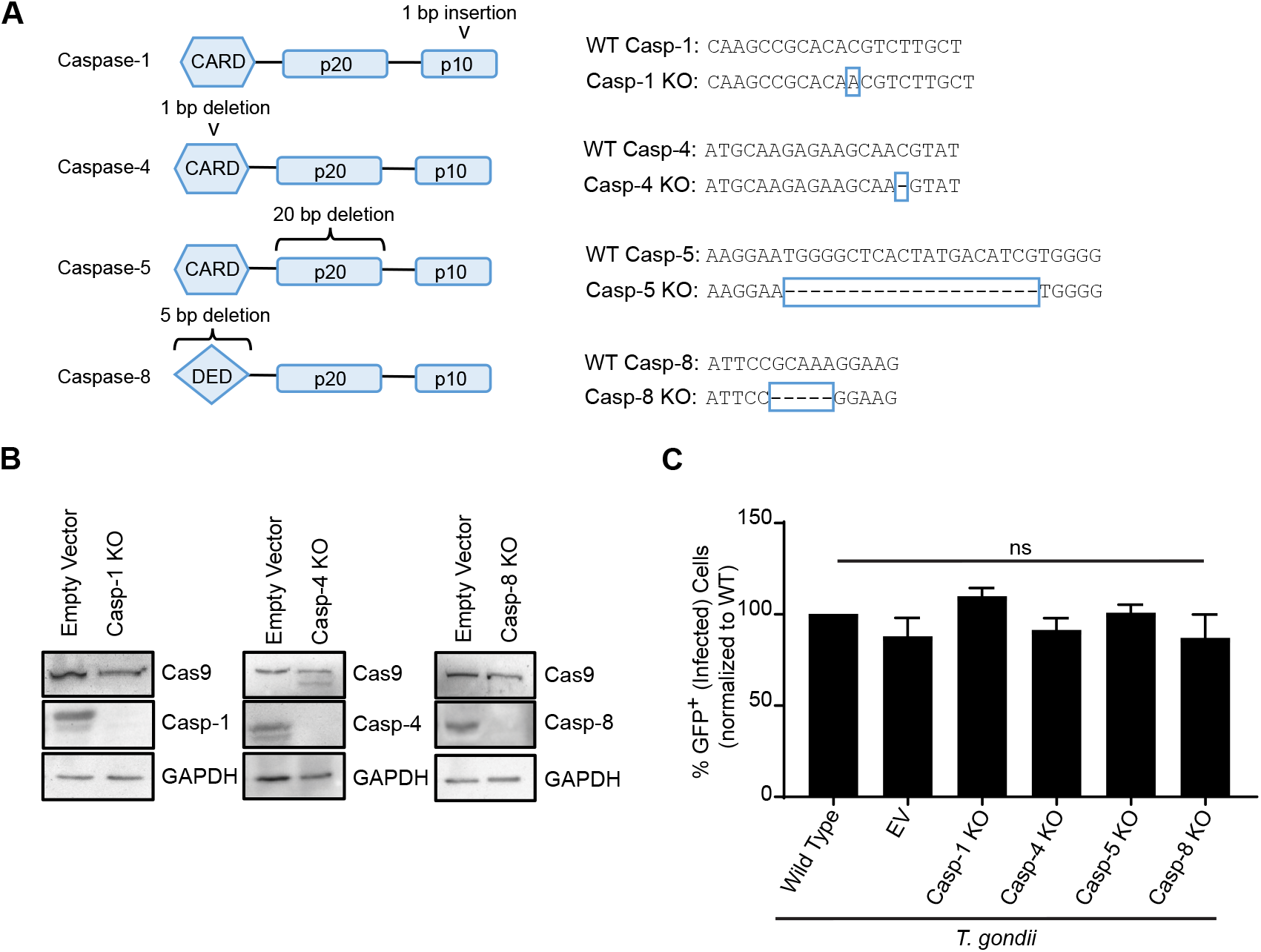
CRISPR-mediated generation and characterization of caspase KO THP-1 cells. **(A)** Schematic details the location of indels in caspase-1, −4, −5, and −8 KO cells. **(B)** EV and caspase-1, −4 and −8 KO THP-1 cells were lysed and probed for Cas9, GAPDH, caspase-1, −4, and −8 by Western blot. **(C)** WT, EV and caspase-1, −4 and −8 KO THP-1 cells were infected with GFP-expressing *T. gondii* at a MOI of 2. GFP^+^ (infected) cells were quantified by flow cytometry. n = 3 experiments. Values are expressed as the mean ± SD, ns (not significant), one-way ANOVA followed by a Tukey post-test.

**Fig. S2.**
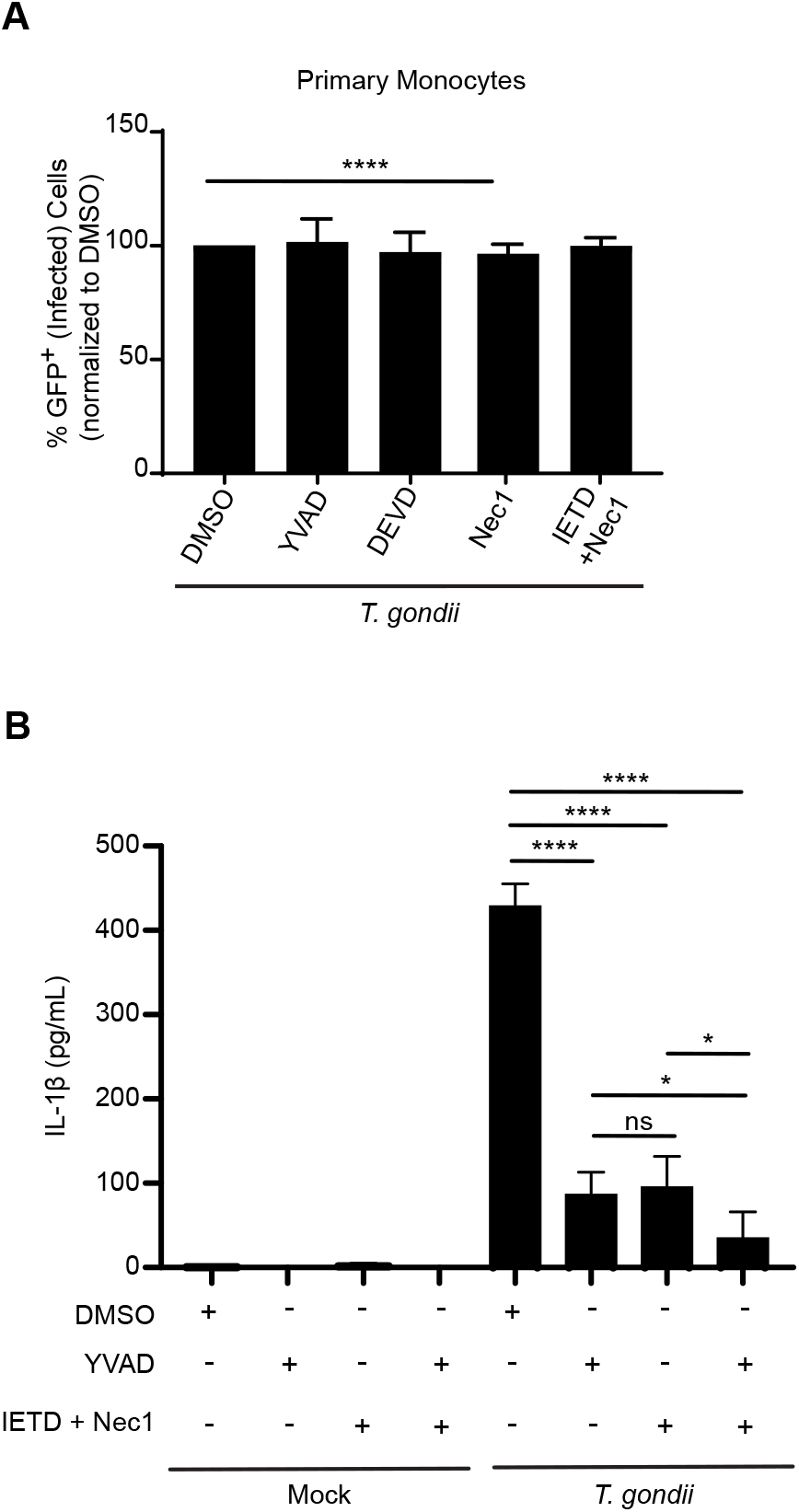
Infection efficiency of caspase and necroptosis inhibitor-treated primary monocytes. Primary human monocytes were pretreated with YVAD, DEVD, Nec1, IETD and Nec1, or an equal volume of DMSO and infected with *T. gondii* at MOI of 2. Infection efficiency, measured by the percentage of GFP^+^ cells, was quantified by flow cytometry. n = 3 experiments. Values are expressed as the mean ± SD, (one-way ANOVA followed by a Tukey post-test). **(B)** EV THP-1 cells were pretreated with DMSO, YVAD, Nec1 and IETD, or combined YVAD, Nec1, and IETD. They were then mock treated or infected with *T. gondii* at an MOI of 2. IL-1β in the supernatant was measured by ELISA. n = 4 experiments. Values are expressed as the mean ± SD, ns (not significant), * *P* < 0.05, **** *P* < 0.0001 (one-way ANOVA followed by a Tukey post-test).

**Fig. S3.**
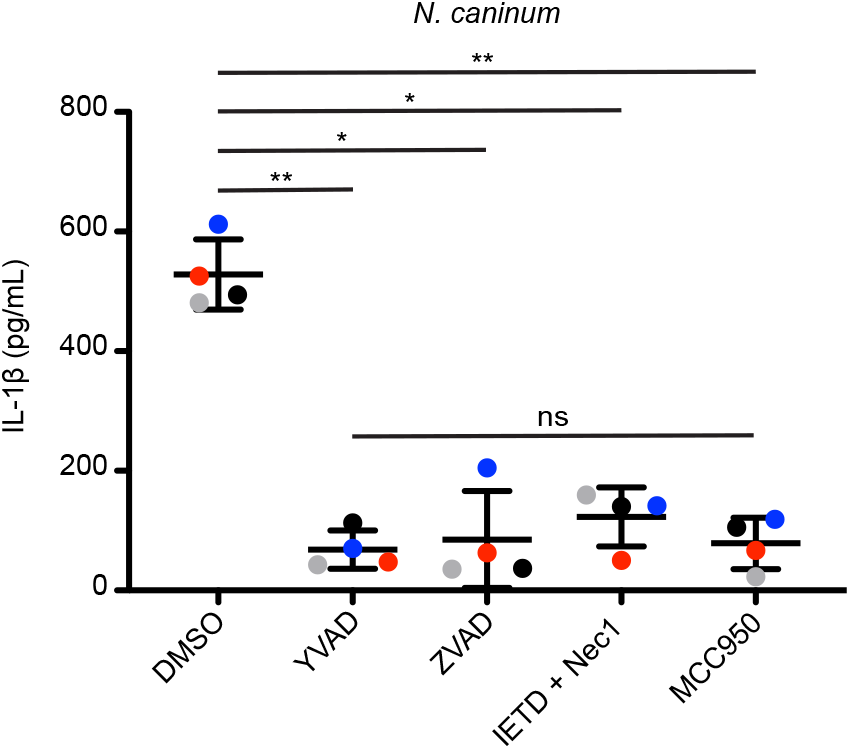
*Neospora caninum*-induced IL-1β release from primary human monocytes involves caspase-1, caspase-8, and the NLRP3 inflammasome. Primary human monocytes were pretreated with YVAD, ZVAD, IETD and Nec1, MCC950, or an equal volume of DMSO. Cells were then mock treated or infected with *N. caninum* at MOI of 2 for 4 hr. IL-1β in the supernatant was measured by ELISA. Each colored dot represents a separate human blood donor. n = 4 experiments with independent blood donors. Values are expressed as the mean ± SD, * *P* < 0.05 (one-way ANOVA followed by a Tukey post-test).

## Notes

### Competing Interest Statement

The authors have declared no competing interest.

